# Effects of Cholesterol on Nanodisc Formation and Magnetic Alignment in DMPC and Glycyrrhizic Acid Systems Probed by ^31^P and ^14^N Solid-State NMR

**DOI:** 10.64898/2026.08.26.747314

**Authors:** Md Rokonujjaman, Sungsool Wi, Ayyalusamy Ramamoorthy

## Abstract

Nanodiscs and bicelles are widely used as membrane mimetics for structural studies of membrane-associated systems. Studies have reported that their magnetic alignment behavior and phase stability are highly sensitive to composition and temperature. In this study, we systematically investigate the effects of cholesterol on bicelle formation and magnetic alignment in DMPC + 0.2 glycyrrhizic acid (GA) systems using a combined ^31^P and ^14^N solid-state NMR experimental and simulation-based approach. Temperature dependent ^31^P NMR spectra reveal a clear transition from vesicle dominant to aligned bicelles/nanodsics phase, while ^14^N quadrupolar splitting and lineshape analysis provides quantitative insights into heterogeneous lipid bilayer populations, distinguishing large aligned nanodiscs (B(L)), small nanodiscs (B(S)), and isotropic/random components (B(R)). A strong correlation is observed between the ^31^P derived bicelle fraction and the ^14^N B(L) population, confirming that macroscopic alignment in the presence of an external magnetic field directly reflects the growth of large, well-ordered nanodiscs. Cholesterol is found to play a critical dual role by modulating membrane order and curvature. At low cholesterol concentration (0-5 mol%), nanodiscs alignment occurs gradually with increasing temperature, while at higher cholesterol concentration (15–25 mol%), the alignment is delayed and accompanied by broader spectral features, indicating structural heterogeneity. Notably, 10 mol% cholesterol consistently provides the optimal balance, enabling efficient temperature dependent conversion to aligned bicelles while maintaining high B(L) populations (∼70-80%) and minimal isotropic fractions. In contrast, higher cholesterol maintains significant B(S) and B(R) populations, even at elevated temperature. The ^14^N quadrupolar coupling (C_q_≈8.5–9.2 kHz for aligned nanodiscs) remains nearly invariant across compositions, showing that cholesterol does not change local headgroup dynamics but instead redistributes lipid populations. These findings establish a combined ^31^P and ^14^N solid -state NMR approach provides a valuable platform for quantitatively correlating membrane structure, dynamics, and alignment, offering practical guidelines for optimizing bicelle systems for high resolution solid-state NMR studies of membrane associated biomolecules.

## Introduction

Biological membranes consist of lipids, proteins, and sterols, which collectively determine cellular structure and function. The fluid-mosaic model initially characterized membranes as dynamic systems with mobile components, providing a foundational understanding of membrane organization.^1^ Subsequent studies revealed that membranes contain small, specialized domains, such as lipid rafts enriched in cholesterol and sphingolipids, which regulate signaling, transport, and structural organization.^2–5^ The physicochemical properties of lipid bilayers, including thickness, morphology, flexibility, and phase behavior, are dictated by lipid composition and hydration levels.^6,7^ Cholesterol plays a critical role by modulating lipid packing, promoting the formation of lipid-ordered phases, and regulating membrane fluidity and stability.^8–14^ These functions are essential for maintaining membrane integrity and mediating protein-lipid interactions, which are fundamental to biological activity.^10–14^

To investigate these complex systems under controlled conditions, researchers have developed a variety of model membrane platforms. Bicelles or nanodiscs, which are composed of long-chain phospholipids such as DMPC and short-chain lipids or amphiphiles, have emerged as versatile membrane mimetics because they can form discoidal nanoparticles suitable for solution NMR based studies and large size structures that magnetically align and enable solid-state NMR applications.^9,15–20^ The formation of bicelles/nanodiscs depends on a balance among bilayer elasticity, edge energy, and amphiphile partitioning, resulting in structures with a planar lipid region and a curved rim.^17–25^ In DMPC-based systems, amphiphilic molecules such as saponins provide an alternative to conventional detergents (such as DHPC) for stabilizing bicelle-like assemblies. While the term “bicelle” was used for the lipid + detergent system, the introduction of MSP based “nanodisc”^26,27^ has expanded the list of belt molecules (including peptides, polymers, and saponins) and the scope of these membrane mimetics; therefore, here onwards, we will use “nanodisc” instead of “bicelle”. Glycyrrhizic acid (GA), a triterpenoid saponin with amphiphilic properties, can insert into lipid bilayers and preferentially localize at the nanodisc rim, reducing edge energy and stabilizing discoidal structures.^28–35^ This process enables the formation of DMPC + GA nanodiscs, which serve as nonionic membrane mimetics with tunable size, curvature, and alignment characteristics.^28–30^ These systems offer a valuable platform for structural studies while maintaining bilayer-like environments suitable for membrane-associated biomolecules.

Cholesterol is a key determinant of membrane structure and phase behavior, and these effects extend to bicelle systems. Cholesterol enhances lipid packing, increases bilayer thickness, and promotes ordering of fatty acid chains, thereby stabilizing liquid-ordered phases and modulating membrane flexibility.^36–38^ In bicelles, these modifications influence both morphology and magnetic alignment by altering the balance between bilayer rigidity and edge-induced stress.^36–39^ Temperature further modulates these processes by dictating lipid phase transitions and facilitating the structural rearrangements necessary for bicelle formation.^37–40^ As temperature increases, lipid bilayers transition from a gel to a liquid-crystalline state, promoting bicelle alignment; however, cholesterol can modify these transitions by maintaining membrane order and impeding structural reorganization.^41–44^ At moderate concentrations, cholesterol enhances bicelle stability and alignment, whereas higher concentrations induce greater structural heterogeneity and obscure the transition from vesicles to bicelles.^43–53^ Glycyrrhizic acid (GA) also influences this equilibrium by interacting with both lipids and cholesterol, thereby altering membrane morphology, permeability, and domain organization.^39^ Collectively, these factors establish a complex interplay between composition and temperature that governs bicelle formation and alignment.

Solid-state NMR spectroscopy provides a robust approach for investigating membrane structure across multiple length scales. ^31^P NMR is particularly valuable due to its high sensitivity to the orientation of phospholipid headgroups, enabling direct assessment of overall membrane organization.^28,30,54–58^ In bicelle systems, ^31^P chemical shift anisotropy (CSA) allows for clear differentiation among isotropic vesicles, lamellar phases, and magnetically aligned bicelles.^28,30,59–63^ In contrast, ^14^N NMR probes the quadrupolar interactions of the choline headgroup, yielding detailed insights into local order, orientation, and the electrostatic environment.^28,64–68^ Because quadrupolar coupling is highly sensitive to molecular motion and alignment, ^14^N NMR serves as a local probe that complements the global information obtained from ^31^P NMR.^69–73^ Recent advancements in solid-state NMR, including enhanced sensitivity, multidimensional experiments, and bicelle-specific methodologies, have significantly improved the resolution and interpretation of membrane structures.^21,25,29,30,74–86^

Concurrently, the simulation of NMR spectra serves as a vital computational bridge between experimental observables and quantitative structural and dynamic information of lipid bilayers and their interactions with peptides.^87,88^ Theoretical lineshape analysis is particularly useful for characterizing anisotropic spectral features associated with dynamic supramolecular assemblies, such as toroidal pores and membrane thinning.^88^ These simulation approaches can also incorporate analyses of 2D exchange spectra and 1D stimulated-echo intensities to extract reorientation-angle distributions, providing insights into membrane morphological changes, including curvature variations, vesicle fusion, and fragmentation.^87^ Furthermore, dynamic NMR lineshape simulation enable the modeling of lateral lipid diffusion, membrane thinning, and other dynamic processes in oriented membrane systems such as bicelles and nanodiscs.^89,90,91^ Together, these methodologies provide a comprehensive understanding of membrane structure, dynamics, and peptide-induced perturbations that cannot be obtained directly from experimental NMR spectra alone.

In this study, we examined how cholesterol concentration and temperature govern the organization and phase behavior of DMPC/glycyrrhizic acid (GA) bicelles. Using ^31^P and ^14^N solid-state NMR spectroscopy, supported by detailed spectral simulations, we tracked cholesterol-dependent changes in bicelle formation, magnetic alignment, and the relative populations of aligned bicelles, small bicelles, and isotropic/vesicular assemblies across a temperature range of 23–60 °C and cholesterol contents from 0 to 25 mol%. By systematically varying cholesterol content and temperature, we establish a framework linking membrane composition to bicelle alignment and structural heterogeneity. This integrated experimental and simulation approach yields new insights into cholesterol-mediated membrane organization in GA-stabilized bicelles and demonstrates the utility of multinuclear solid-state NMR for probing complex membrane systems.

## 2. Experimental and Methods

### 2.1. Materials

1,2-Dimyristoyl-sn-glycero-3-phosphocholine (DMPC, >99%) was obtained from Avanti Polar Lipids (Alabaster, AL, USA), and glycyrrhizic acid (GA) monoammonium salt (>98%) was sourced from Thermo Fisher Scientific (Waltham, MA, USA). All other chemicals were acquired from Sigma-Aldrich (USA) and used without further purification. DMPC served as a model phospholipid and GA as a rim-forming amphiphile, consistent with established bicelle and saponin-based membrane systems in the literature.^9,24,28,30–32^ The molecular structures of DMPC, GA, and cholesterol are presented in Figure S1.

### 2.2. Preparation of DMPC + GA Bicelles and Cholesterol-Containing Samples

DMPC and glycyrrhizic acid (GA) bicelles were prepared with modifications to the protocols described by McCalpin et al. (2026) and De Angelis et al. (2022).^30,50^ DMPC (100 mg/mL) and GA (0.2 w/w; 20 mg/mL) were co-dissolved in chloroform. The solvent was removed under a gentle stream of nitrogen gas, followed by overnight lyophilization to ensure complete solvent removal. The resulting lipid film was hydrated in 10 mM Tris buffer containing 100 mM NaCl at pH 7.4, and GA was added. The mixture underwent 3– 5 freeze–thaw cycles with mixing between cycles. For cholesterol-containing samples, DMPC and cholesterol were co-dissolved in chloroform prior to film formation. Stock solutions of DMPC (200 mg/mL) and cholesterol (50 mg/mL) were prepared in chloroform. Each sample contained 20 mg of DMPC, with cholesterol added in amounts of 0, 0.57, 1.14, 1.71, 2.28, and 2.86 mg, corresponding to DMPC to cholesterol molar ratios of 0, 0.05, 0.10, 0.15, 0.20, and 0.25 (or 0, 5, 10, 15, 20, and 25 mol% cholesterol relative to DMPC). The organic solvent was removed under nitrogen, and the sample was lyophilized overnight. The dried films were hydrated in 200 μL of 10 mM Tris buffer with 100 mM NaCl at pH 7.4, and GA (4 mg) was added to achieve the desired 0.2 GA composition. The mixtures were subjected to 3–5 freeze–warm–freeze cycles, with mixing between cycles to ensure sample uniformity.

### 2.3. ^31^P and ^14^N Solid State NMR Spectroscopy

^31^P NMR experiments were conducted on a 400 MHz Bruker solid-state NMR spectrometer using a low-E static 5 mm double-resonance HX probe built at the National High Magnetic Field Laboratory (NHMFL), USA, with resonance frequencies set to 161.97 MHz for ^31^P and 400.11 MHz for ^1^H.^92^ Data acquisition employed a 4.2 μs 90° pulse (80 W), a 2.0 s recycle delay, and a 5.75 μs 90° pulse with 40 W waltz64 proton decoupling.^93^ Liquid H_3_PO_4_ served as the reference to set the chemical shift to 0 in the ^31^P NMR spectra. NMR signals were averaged over 512 scans and processed with Gaussian broadening (LB = −20; GB = 0.02). ^14^N NMR experiments were carried out by tuning the X channel of the HX probe to 28.93 MHz for ^14^N. Data were collected using a 90° – tau – 90° solid-echo pulse sequence. A 7.5 μs 90° pulse (120 W), a 1.0 s recycle delay, and a 5.25 μs 90° pulse with 80 W waltz64 proton decoupling were used for the experiments. Liquid NH_4_Cl served as the reference to set the chemical shift to 0 in the ^14^N NMR spectra. NMR signals were averaging over 10,000 scans and processed with Gaussian broadening (LB = −20; GB = 0.02). For both ^31^P and ^14^N NMR experiments, samples were equilibrated for 30 minutes at each temperature prior to data collection.

### 2.4. Spectral simulations

Anisotropic ^31^P and ^14^N solid-state NMR spectra provide structural and dynamical information on lipid assemblies through orientation-dependent chemical shift anisotropy (CSA) and quadrupolar (QC) interactions.^87,88,94–96^ In hydrated membranes, rapid uniaxial rotation of lipid molecules about their long axis leads to motional averaging of these interactions, yielding effectively axially symmetric tensors.^67^ The extent of this averaging is described by an order parameter that reflects lipid mobility. The observed resonance frequencies depend on the orientation of these motionally averaged tensors relative to the external magnetic field. For bicelles, the lipid bilayer normal defines the reference axis, with native magnetic alignment typically corresponding to a perpendicular orientation relative to the field (*n* ⊥ *B*_0_); alternative alignments can be induced using paramagnetic additives.^71,97^ In curved regions of standard DMPC:DHPC bicelles occupied by DHPC molecules, the resulting ssNMR lineshapes arise from a distribution of local orientations across the membrane surface. Our recently developed lipid simulation protocol reproduces these lineshapes by integrating over orientation space using geometry-dependent probability distributions that reflect the underlying membrane morphology.^89^ The simulation framework accounts for bicelle geometries comprising planar bilayer regions and a curved rim, defined by parameters such as bilayer thickness and the q-factor governing lipid partitioning.

As the rim region of the DMPC:GA nanodiscs is occupied by GA, which lacks both ^31^P and ^14^N nuclei, the spectral simulations were performed exclusively for the flat region of the nanodiscs. Consequently, lipid lateral diffusion was not incorporated into the simulations, as it does not affect the spectra of lipids in the flat region when the nanodisc normal is oriented either parallel or perpendicular to the external magnetic field, B_0_.Because the bicelles contributing to the NMR signals from the flat disc regions adopt a well-defined alignment with *n* ⊥ *B*_0_, and we consider motionally averaged uniaxial CSA (^31^P) or quadrupolar (^14^N) tensors (η = 0), the simulation employed a single effective orientation. In addition, a random vesicle model combined with a diffusive jump approach was used to reproduce fully or partially averaged isotropic lineshapes. Vesicles exhibit an isotropic orientational distribution; to represent this randomness, a spherical geometry was adopted in the simulation, and a powder average over 2000 orientations was used. All simulations were implemented using MATLAB code.^89^ Spectra were generated by numerically sampling orientations, propagating magnetization, and Fourier transforming the resulting signals. To account for experimental linewidths, Lorentzian broadening (50– 800 Hz) or Gaussian broadening (LB = −20 to −30 Hz; GB = 0.02–0.1) was applied.

## 3. Results

### 3.1. ^31^P Solid-State NMR Confirms DMPC + GA Bicelle Formation

One-dimensional static ^31^P solid-state NMR spectroscopy was employed to investigate the self-assembly and magnetic alignment of DMPC and glycyrrhizic acid (GA) mixtures as a function of temperature. The ^31^P NMR signal is highly sensitive to lipid organization, as the phosphate chemical shift anisotropy (CSA) reflects the orientation of phospholipid headgroups relative to the magnetic field.^19,28,30,98–104^ In lamellar systems such as multilamellar vesicles (MLVs), randomly oriented bilayers produce a characteristic powder-pattern lineshape, whereas magnetically aligned bicelles exhibit a narrow resonance due to uniaxial alignment of lipid molecules and motional averaging.^21,30, 74,105–108^ As shown in Figure 1A, pure DMPC (12% w/v) forms multilamellar vesicles with a classic powder-pattern spectrum, which arises from randomly oriented lipid molecules, remains largely unchanged with temperature, consistent with established behavior for phosphatidylcholine membranes in the lamellar phase.^19,30,74, 109–115^ To further validate this behavior, simulated ^31^P spectra of DMPC MLVs are presented in Figure S2, generated at two representative temperatures: 23 °C, below the gel-to-liquid-crystalline phase transition of DMPC (∼24 °C), and 38 °C, above it. In both cases, the simulations reproduce the characteristic powder-pattern lineshape, confirming that the axially symmetric CSA tensor and overall bilayer organization are preserved across the phase transition. Upon the addition of GA, significant changes in lipid organization were observed, indicating disruption of lamellar packing and the formation of non-lamellar structures. In the DMPC + 0.15 GA system (Figure 1B), shows powder-pattern shape spectrum at 23 °C. As the temperature increases (≥ 29 °C), the broad powder-pattern transitions to a narrow resonance near −10 ppm, characteristic of magnetically aligned bicelles with the lipid bilayer normal oriented perpendicular to the magnetic field axis.^21,30,98^ This transition indicates the formation of lipid assemblies with lipids rotating rapidly around the bilayer normal, averaging the CSA and producing a sharp, isotropic-like signal.^19,21,30,55,98^

**Figure 1.**
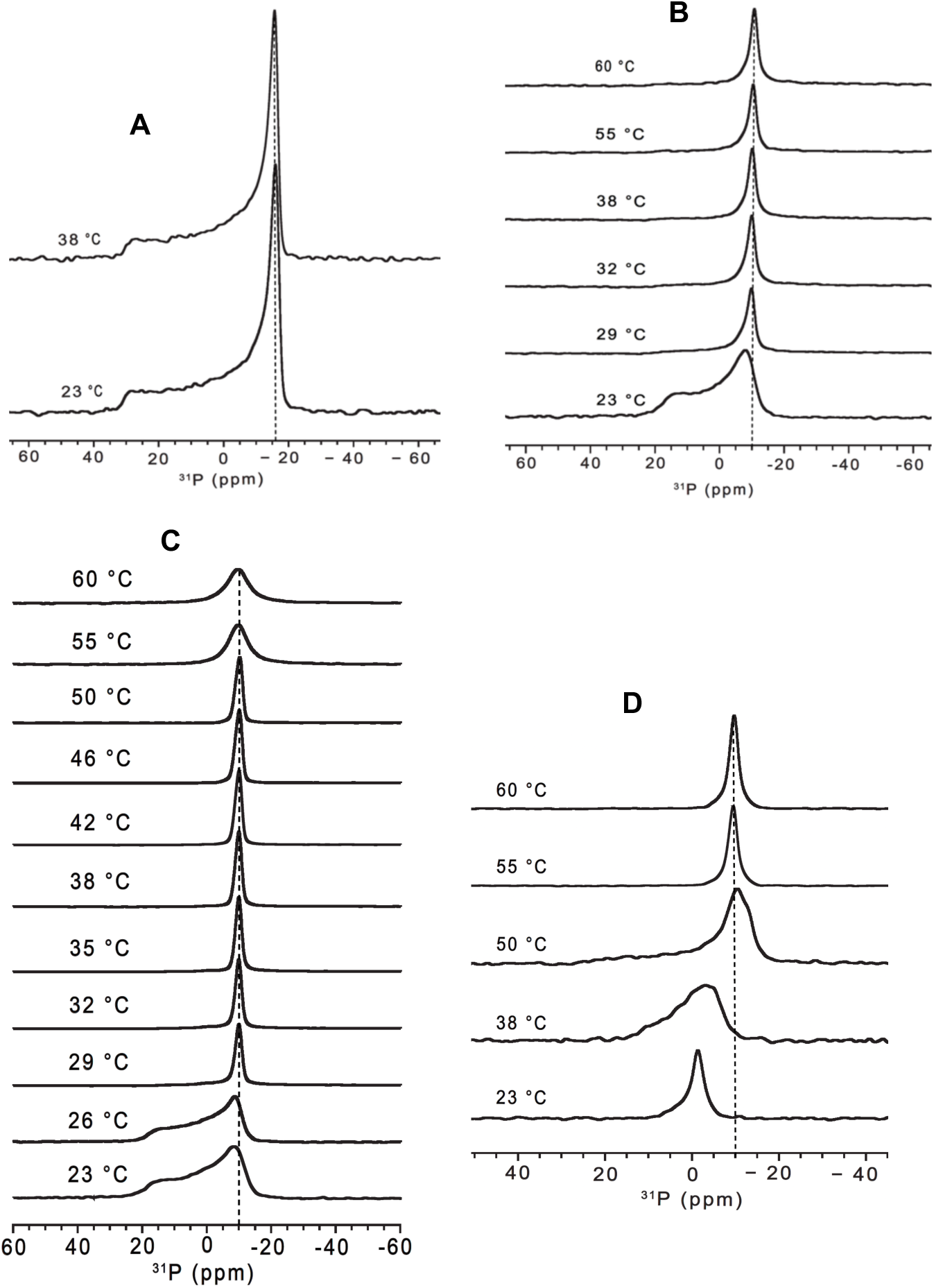
Static ^31^P solid-state NMR spectra were acquired at increasing temperatures in 10 mM Tris buffer containing 100 mM NaCl at pH 7.4 for the following samples. (A) DMPC multilamellar vesicles (MLVs, 12% w/v), which display a characteristic axially symmetric powder pattern lineshape. (B) DMPC (100 mg/mL) with 0.15 GA (15 mg/mL) bicelles, where the onset of magnetic alignment is observed at temperatures of 29 °C or higher. (C) DMPC (100 mg/mL) with 0.20 GA (20 mg/mL) bicelles, which demonstrate well-defined and stable alignment beginning at approximately 29 °C. (D) DMPC (100 mg/mL) with 0.25 GA (25 mg/mL) bicelles, where alignment is delayed and occurs only at elevated (figure 1 contn.) temperatures of 55 °C or higher. The dashed line at ppm marks the characteristic resonance position of magnetically aligned bicelles/perpendicular edge, as observed in the spectra.

The DMPC + 0.2 GA system (Figure 1C) demonstrates the most efficient bicelle formation exhibiting a sharp, symmetric resonance at −10 ppm, for temperatures above the main gel to liquid crystalline phase transition temperature of DMPC (> 24 °C), from 29 °C to at least 60 °C. These results indicate that bicelle formation is highly cooperative and that magnetic alignment is stable across a broad temperature range. This efficient alignment is consistent with previous studies demonstrating that amphiphilic additives and saponin-like molecules stabilize discoidal bicelles by reducing edge energy.^30,50,72,84,98,116^ The disappearance of powder-pattern edges and the emergence of a sharp resonance confirm the formation of a uniformly aligned bicelle phase. This structural reorganization reflects a transition from randomly oriented vesicles to magnetically aligned discoidal assemblies. GA acts as an effective amphiphilic modulator, forming the bicelle rim, where it reduces edge energy and stabilizes the flat bilayer region, thereby promoting bicelle formation^.30,39,50,56,89,98^

At higher GA concentrations (0.25 GA, Figure 1D), the spectra reveal increased structural heterogeneity. At 23 °C, a sharp isotropic resonance near 0 ppm is observed, consistent with small, rapidly tumbling vesicles or micelle-like aggregates. At 35 °C, the resonance broadens and becomes less symmetric, indicating reduced isotropic motion and the onset of bicelle formation. At elevated temperatures (≥50 °C), a main peak near −10 ppm emerges, indicative of aligned bicelles, although a weak powder pattern-like component persists, suggesting the presence of non-aligned domains. Complete alignment is achieved only at 55–60 °C, implying that excess GA increases structural diversity and diminishes alignment cooperativity. In summary, GA concentration strongly influences bicelle formation and alignment. Bicelle alignment initiates at 0.15 GA at temperatures above 29 °C, but the 0.2 GA mixture yields the most rapid, cooperative, and stable alignment. This composition was selected for subsequent cholesterol-dependent studies because it allows higher cholesterol incorporation while maintaining an optimal bicelle q ratio (DMPC/GA molar ratio) and robust magnetic alignment. ^30,50,98,116,117^

### 3.2. ^31^P Solid-State NMR Analysis of Cholesterol-Dependent Modulation of Bicelle Alignment

The static ^31^P solid-state NMR spectra demonstrate that cholesterol significantly influences the temperature-dependent transition from vesicles to bicelles in DMPC + 0.2 GA bicelles (Figure 2). Across all samples, the spectra shift from broad, vesicle-like or non-aligned bilayer line shapes at lower temperatures to a narrow resonance near −10 ppm, characteristic of magnetically aligned bicelles. This observation is consistent with the established sensitivity of ^31^P NMR to phospholipid headgroup orientation and bicelle alignment, as well as with previous studies on magnetically alignable bicellar systems and GA-containing bicelles.^19,21,22,24, 30,39,98,100,106^ For panels A to E (right panel experimental and left panel simulated spectrum), spectral simulations employed a constant ^31^P chemical shift anisotropy (CSA) of approximately 20 ppm with η = 0, indicating axial symmetry. Notably, only the 25 mol% cholesterol sample required an additional bicelle environment with distinct CSAs (20 ppm and 30 ppm), reflecting increased structural diversity. For 5–25 mol% cholesterol, the DMPC + 0.2 GA bicelle samples exhibited either heterogeneous populations of multilamellar vesicles (MLVs) or a mixture of MLVs and aligned bicelles, in contrast to the 0 mol% cholesterol sample. The observed ^31^P powder-pattern contribution arises from the vesicle population and is strongly influenced by the molecular tumbling rate (1/τ_c_, where τ_c_ is the correlation time). As illustrated by the simulated spectra in Figure S3, decreasing τ_c_ (increasing tumbling rate) progressively averages the ^31^P chemical shift anisotropy (CSA), resulting in the continuous transformation of the broad MLV powder pattern into a sharp isotropic resonance centered at 0 ppm. Moreover, different Lorentzian line broadening (LB, Hz) values were used to optimize the simulated ^31^P spectra to match the experimental line shapes. The effect of LB on the spectral line shape is shown in Figure S4.

**Figure 2.**
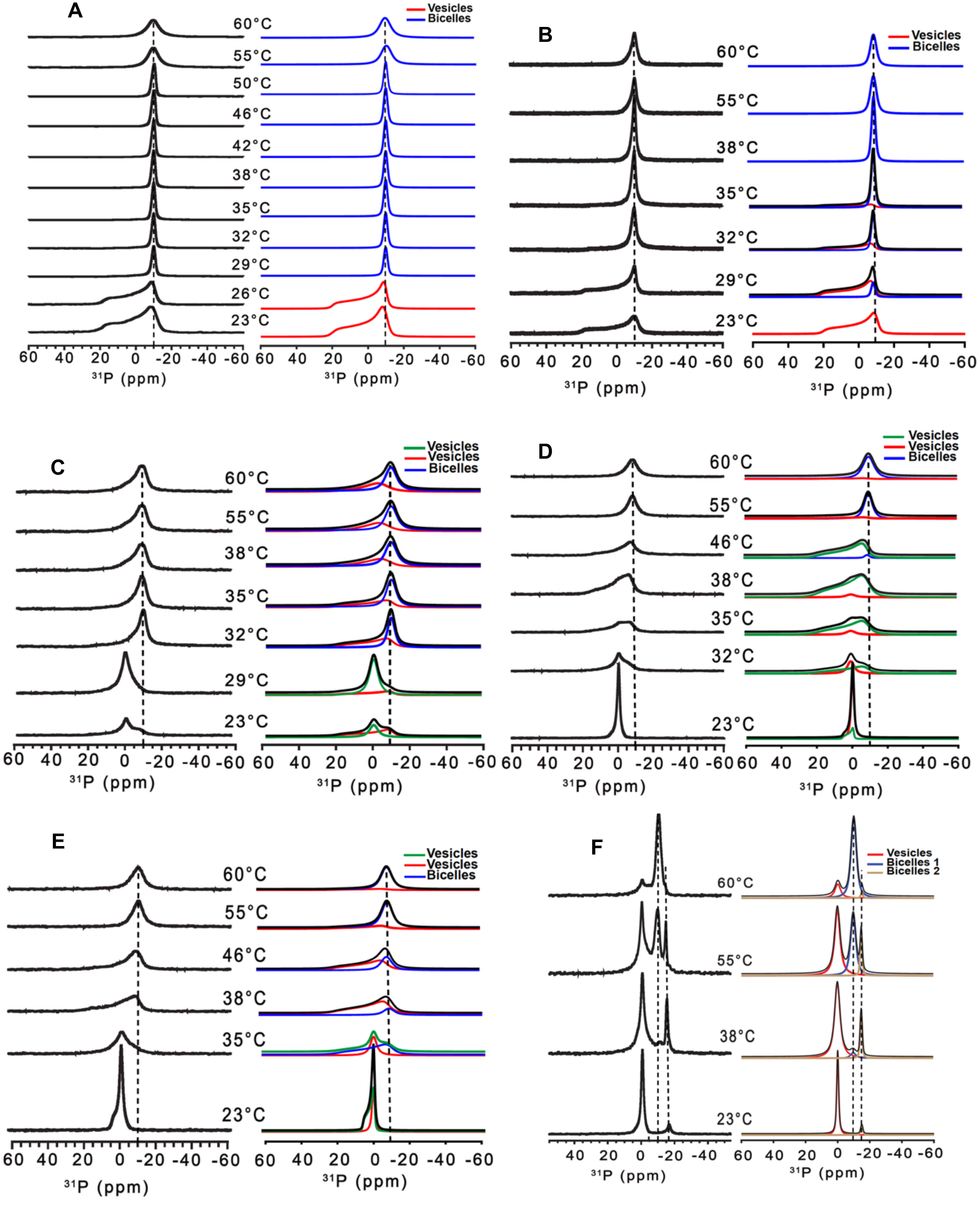
Cholesterol dependent bicelle formation and lipid phase are characterized by static ^31^P solid-state NMR and quantitative spectral simulations, showing experimental (left) and simulated (right) spectra of DMPC (100 mg/mL) + 0.2 GA (20 mg/mL) bicelles in 10 mM Tris buffer with 100 mM NaCl at pH 7.4 as a function of temperature for cholesterol contents of (A) 0, (B) 5, (C) 10, (D) 15, (E) 20, and (F) 25 mol%. For panels A–E, the simulated spectra are well reproduced using a single ^31^P chemical shift anisotropy (CSA) of ∼20 ppm with η = 0. In contrast, panel F (25 mol%) requires two bicelle components with CSA values of ∼20 and ∼30 ppm, consistent with cholesterol-induced structural heterogeneity. All additional simulation parameters for panels A–F are summarized in Table TS1. The dashed line at ∼−10 & −17 ppm marks the characteristic resonance position of magnetically aligned bicelles.

In the cholesterol-free (0 mol%) system (Figure 2A and Table TS1), the spectra exhibit a clear and direct temperature-dependent phase transition. The experimental panel in Figure 2A (left) is identical to Figure 1C, as it is included to present the experimental and simulated spectra side by side and facilitate comparison across different cholesterol mol% samples. At 23–26 °C, the experimental spectra (left panel) display broad, asymmetric powder-pattern line shapes typical of vesicular or non-aligned lipid bilayers. Bicelle alignment initiates at approximately 29 °C, as indicated by spectral narrowing, and at 32 °C or higher, a sharp resonance near −10 ppm emerges, signifying magnetically aligned bicelles. The simulated spectra (right panel) reproduce these features using a CSA of about 20 ppm and η = 0. Line broadening (LB) values of 200– 900 Hz were necessary to match the high-temperature experimental linewidths at 50–60 °C. Collectively, the simulations closely replicate the progressive narrowing and sharpening of the lineshape, confirming a vesicle-to-bicelle transition with improved magnetic alignment as temperature increases. These findings are consistent with earlier ^31^P solid-state NMR studies on DMPC + 0.2 GA bicelles.^30^

At 5 mol% cholesterol (Figure 2B and Table TS1), a similar transition is observed, although the aligned bicelle phase appears at a slightly higher temperature. At 23 °C, the spectrum is fully vesicular (red in the spectra indicating unaligned lipid population), and the simulation corresponds to a 100% vesicle (or unaligned lipids) contribution. At temperatures higher than 23 °C, the spectra exhibit line shapes corresponding to unaligned lipids (termed as vesicles) represented by broad motionally-averaged powder patter and aligned lipids represented by narrow peak around −10 ppm. The percent (%) of lipids population (bicelles or vesicle) present in each lipid phase is determined by simulating the spectra. At 29 °C, the system comprises approximately 15% bicelles and 85% vesicles, and at 32 °C, the bicelle and vesicle populations are nearly equal (approximately 50:50). At 35 °C, the bicelle fraction (blue in the spectra) increases to about 78%, with the remaining 22% as vesicles. Full bicelle alignment is achieved at 38 °C, and above this temperature, the spectra are dominated by the narrow aligned bicelle resonance. The simulated spectra again utilize CSA = 20 ppm and η = 0, with bicelle LB values of approximately 200 Hz at 38 °C and about 400 Hz at 55–60 °C to match the experimental linewidths. Compared with the 0 mol% cholesterol sample, where bicelle alignment begins at approximately 29 °C, the 5 mol% cholesterol sample requires a higher temperature for complete alignment. These results indicate that even a small amount of cholesterol elevates the temperature required for the vesicle-to-bicelle transition. This trend is consistent with the established effect of cholesterol in increasing bilayer packing, order, and membrane rigidity.^38,41,44–48,50,117^

At 10 mol% cholesterol (Figure 2C and Table TS1), the ^31^P spectra reveal a temperature-dependent vesicle-to-bicelle transition with pronounced phase coexistence over a broad temperature range. The experimental spectra are modeled using one bicelle component (blue) and two vesicle components (red and green), with vesicles characterized by correlation times, τc. At 23 °C, the spectrum is dominated by vesicles with τ_c_ = 300 ms and 20,000 ms, indicating slow, heterogeneous dynamics. At 29 °C, the system remains predominantly vesicular (vesicle:vesicle approximately 80:20) with τ_c_ = 300 ms and 13,000 ms. At 32 °C, bicelle formation commences, with a roughly 50:50 bicelle:vesicle population and τ_c_ = 8,000 ms for the vesicle components. From 35 to 60 °C, the bicelle fraction remains around 60%, while vesicles constitute about 40%, with τ_c_ decreasing from 7,000 ms to 1,300 ms, reflecting accelerated motion at elevated temperatures. The bicelle component required LB values of approximately 500–900 Hz, while the vesicles required LB values of about 200–600 Hz. Relative to the 0 mol% system, which transitions more directly to a predominantly aligned bicelle phase, the 10 mol% sample maintains significant vesicle-bicelle coexistence over a wider temperature range. In contrast to the 5 mol% system, which achieves full bicelle alignment at approximately 38 °C, the 10 mol% sample does not reach complete alignment even at higher temperatures. Thus, 10 mol% cholesterol broadens the transition region by stabilizing both vesicle and bicelle populations, while maintaining the phosphate CSA at 20 ppm.

At 15 mol% cholesterol (Figure 2D and Table TS1), the spectra exhibit a more delayed and strongly temperature-dependent phase transition. The simulations incorporate two vesicle components (red and green in the spectra) and one bicelle component (blue in the spectra). At 23 °C, the spectrum is dominated by vesicles (vesicle:vesicle approximately 80:20) with τ_c_ around 200 ms for vesicle 1 and τc around 8,000 ms for vesicle 2. At 32 °C, vesicles remain dominant (vesicle:vesicle approximately 35:65) with τc around 200 ms and 4,000 ms. At 35 °C, the system remains vesicular (vesicle:vesicle approximately 10:90) with τ_c_ around 200 ms and 4,000 ms, and at 38 °C, the vesicle population is still dominant (vesicle:vesicle approximately 5:95) with the same τ_c_ values, indicating persistent vesicle phases. A distinct bicelle contribution first appears at 46 °C, with vesicle:bicelle approximately 95:5 and vesicle τ_c_ around 4,000 ms. At 55 °C, bicelles predominate (vesicle:bicelle approximately 10:90), and at 60 °C, the aligned bicelle fraction increases further (vesicle:bicelle approximately 5:95), while the vesicle τ_c_ remains around 4,000 ms. The bicelle component required LB values of approximately 300 Hz at 46 °C, about 600 Hz at 55 °C, and about 900 Hz at 60 °C, while the vesicle component required LB values of about 200–500 Hz. Compared to 0 mol% cholesterol, the transition is notably delayed; compared to 5 mol%, bicelle alignment becomes dominant only at higher temperatures; and compared to 10 mol%, bicelle formation initiates later, and vesicle phases persist longer. These findings indicate that increasing cholesterol to 15 mol% enhances bilayer packing and rigidity, stabilizing vesicles at intermediate temperatures and necessitating higher temperatures for reorganization into aligned bicelles.

At 20 mol% cholesterol (Figure 2E and Table TS1), the spectra also display a delayed and mixed vesicle-to-bicelle transition. The simulations again employ one bicelle component and two vesicle components. At 23 °C, the spectrum is dominated by vesicles with two populations (vesicle:vesicle approximately 40:60) with τc around 200 ms and 4,000 ms. At 35 °C, the system remains vesicular (vesicle:vesicle approximately 35:65), with τc values of about 200 ms and 4,000 ms, indicating persistent vesicle phases. At 38 °C, the system is still predominantly vesicular (vesicle:bicelle approximately 75:25) with vesicle τ_c_ around 4,000 ms. At 46 °C, bicelle formation becomes significant (vesicle:bicelle approximately 60:40), while vesicle dynamics remain at τ_c_ around 4,000 ms. At 55 °C, bicelles dominate (vesicle:bicelle approximately 20:80), and at 60 °C, strong alignment is observed (vesicle:bicelle approximately 10:90). The bicelle component required LB values of approximately 800 Hz from 38 to 60 °C, while vesicle components required LB values of about 200–500 Hz. Compared to samples with lower cholesterol, the 20 mol% system exhibits a broader, more heterogeneous transition. Relative to 15 mol%, there is a minor difference, as bicelle formation begins at 38 °C for 20 mol% but at 46 °C for 15 mol%. Nevertheless, the overall trend is evident: from 0 to 20 mol% cholesterol, increasing cholesterol progressively stabilizes vesicular structures over a wider temperature range and requires higher temperatures for reorganization into aligned bicelles, while the phosphate CSA remains approximately 20 ppm. This is consistent with the established effects of cholesterol on membrane order, thickness, and rigidity.^45,46,48.50.52^

The 25 mol% cholesterol sample (Figure 2F and Table TS1) exhibits the most heterogeneous behavior. In this case, the experimental spectra required three simulated components: one vesicle component and two distinct bicelle components (B1 and B2). The vesicle component was modeled with CSA = 20 ppm and τ_c_ = 50 ms at 23 °C and τ_c_ = 140 ms at 38–60 °C. Since vesicle tumbling rate = 1/ τ_c_, the higher τc at elevated temperature indicates slower vesicle motion in the model. Bicelle 1 (B1) was assigned CSA = 20 ppm and LB = 400 Hz, while bicelle 2 (B2) required CSA = 30 ppm and LB = 100 Hz. The vesicle component used LB = 70 Hz at 23 °C and LB = 300 Hz at 38–60 °C. At 23 °C, the spectrum is dominated by vesicles, with a minor contribution from bicelle 2 (∼17 ppm), resulting in V:B2 = 90:10. At 38 °C, the system remains predominantly vesicular, with V:B1:B2 = 79:5:16. At 55 °C, all three components contribute substantially, with V:B1:B2 = 50:40:10, indicating strong coexistence of vesicles and two bicelle environments. At 60 °C, the spectrum is primarily bicellar, with V:B1:B2 = 16:82:2, indicating that bicelle 1 at approximately at −10 ppm is the principal aligned phase at high temperature, while vesicles and bicelle 2 are minor. Thus, bicelle formation persists at 25 mol% cholesterol at high temperature, but the transition is considerably more heterogeneous than in the 0, 5, 10, 15, and 20 mol% systems. The requirement for two bicelle populations suggests that very high cholesterol concentrations favor multiple aligned bicelle environments rather than a single one.

Across the entire cholesterol range from 0 to 25 mol%, the ^31^P spectra clearly demonstrate that cholesterol modulates bicelle formation, alignment, and phase coexistence. The 0 mol% sample exhibits the most direct transition to a single aligned bicelle phase. At 5 mol%, alignment remains robust but occurs at a slightly higher temperature. At 10 mol%, bicelle formation initiates early, at approximately 32 °C, and the system maintains a high, stable bicelle fraction, although vesicles persist over a broader temperature range. At 15 and 20 mol%, the transition is delayed and extended, with vesicles remaining dominant over a wider temperature range and full alignment requiring higher temperatures. At 25 mol%, the system is the most heterogeneous, necessitating multiple bicelle populations for accurate modeling. In summary, increasing cholesterol content enhances bilayer packing and rigidity, thereby delaying the phase transition, broadening the coexistence range, and increasing structural diversity. In contrast, 10 mol% cholesterol gives the best balance between flexibility and order in the membrane, which helps bicelles form and align efficiently. At this level, the membrane is stable enough to support large bicelles but still flexible enough to allow easy structural rearrangement. As a result, more aligned bicelles are formed, fewer disordered structures remain, and the alignment stays stable over a wide temperature range.^30,38,39.45,46,48,50,52,98,116^

### 3.3. ^14^N Solid-State NMR Reveals Cholesterol-Dependent Bicelle Population Redistribution

The ^14^N solid-state NMR spectra provide direct information on lipid headgroup ordering and enable measurement of bicelle populations via quadrupolar lineshape analysis. While ^31^P NMR shows global magnetic alignment, ^14^N NMR focuses on the local electrostatic environment, orientation and motion of the choline headgroup. This allows distinguishing between aligned bicelles, small bicelles, and isotropic components.^28,25,41,56,57,59,60,89,118.119^ In multilamellar vesicles in which lipids are randomly oriented, a motionally-averaged ^14^N quadrupolar powder pattern is observed. The near tetrahedral symmetry around ^14^N reduces the quadrupole coupling, in addition to the axial motion of lipids. In aligned systems, the uniaxial alignment of lipid molecules and therefore the choline group result in narrower spectral lines at the perpendicular edges of the ^14^N quadrupole coupling powder pattern.

For pure DMPC vesicles (Figure S5), the spectra show a typical powder-pattern line shape with C_q_ around 14.5 kHz, which matches randomly oriented bilayers.^25,56,57,59,60,89^ Raising the temperature only slightly affect the powder-pattern and does not change C_q_, indicating that the local dynamic environment of the headgroup remains unchanged.^59,60^ Moreover, molecular tumbling strongly influences the ^14^N quadrupolar line shape, as demonstrated by the simulated spectra in Figure S6. At long correlation times (τ_c_ = 10,000–4000 ms), the spectra exhibit the characteristic quadrupolar powder pattern of slowly tumbling phospholipid vesicles, indicating negligible motional averaging. As τ_c_ decreases (increasing tumbling rate), the quadrupolar interaction is progressively averaged, leading to the collapse of the powder pattern into a narrow isotropic resonance at 0 kHz in the fast-motion regime (τ_c_ = 100–10 ms), characteristic of rapidly tumbling membrane assemblies such as small vesicles or micelles. The quadrupolar coupling constant (C_q_) strongly influences the ^14^N quadrupolar lineshape, as demonstrated by the simulated spectra in Figure S7. Increasing C_q_ from 0.5 to 12.5 kHz progressively increases the quadrupolar splitting and broadens the powder pattern, whereas decreasing C_q_ collapses the spectrum toward a narrow resonance centered at 0 kHz. In contrast DMPC MLV’s, DMPC + 0.2 GA bicelles exhibit partially averaged quadrupolar interactions due to solubilization of vesicles by GA and fast rotational diffusion.^28,57,72,105, 121,122,123^

The spectra were analyzed using a three-component model: large aligned bicelles (B(L)) exhibiting large quadrupolar splitting, small bicelles (B(S)) exhibiting smaller quadrupolar splitting, and isotropic or random assemblies (B(R)) represented by the isotropic peak at 0 ppm, with an axially symmetric electric field gradient (η = 0). The aligned bicelle component always has δ_iso_ about 27 ppm and C_q_ between 8.5 and 9.2 kHz, while the isotropic or small bicelle component appears near δ_iso_ about 0 ppm with a lower C_q_ of 0.5 to 1.0 kHz. These values remain nearly the same across different temperatures and cholesterol levels, indicating that cholesterol does not alter the basic headgroup electronic environment but instead affects phase of lipid populations. The evolution of lipid populations in the DMPC + 0.2 GA bicelle system was investigated by simulating the ^14^N NMR spectra with varying relative populations of large bicelles (B_L_), small bicelles (B_S_), and randomly aggregated species including vesicles (B_R_) (Figure S8). Increasing the B_R_ population progressively decreases the intensity of the aligned bicelle resonance at δ_iso_ ≈ 27 ppm while enhancing the isotropic component near 0 ppm, reflecting the transition from well-aligned bicelles to randomly oriented membrane assemblies. The simulated spectra reproduce the coexistence of aligned and isotropic components, demonstrating that the experimental ^14^N line shapes arise from mixed lipid populations with different degrees of molecular order and mobility.

In the cholesterol-free system (0 mol%, Figure 3A & Table TS2), the ^14^N analysis shows a clear shift from isotropic assemblies to aligned bicelles as temperature rises. At 23 °C, there are equal amounts of aligned bicelles and isotropic components (B(L):B(R) = 50:50), showing partial ordering even at low temperature. As the temperature goes up to 29 °C and 35 °C, the aligned bicelle population rises to 59% and 72%, while the isotropic fraction drops. At 38 °C, B(L) stays high at 64% and B(S) increases slightly to 17%, suggesting some redistribution into smaller bicelles. At 55 °C, B(L) is still dominant (about 69%) and the isotropic part is lower (about 15%). This pattern matches the ^31^P results, confirming that higher temperature encourages bicelle alignment and reduces isotropic disorder, as expected.^21,24,54,71–74^ At 23 and 29 °C, ^14^N NMR shows a significant aligned bicelle population (B(L) about 50%), which reflects local ordering of the phosphatidylcholine headgroups. In contrast, ^31^P NMR only shows an isotropic vesicular lineshape, meaning there is no long-range magnetic alignment. This difference is because ^14^N NMR is sensitive to local headgroup orientation and can detect partially ordered assemblies, while ^31^P NMR needs uniform, large-scale alignment to show the typical aligned bicelle signal.^25,54,59^

**Figure 3.**
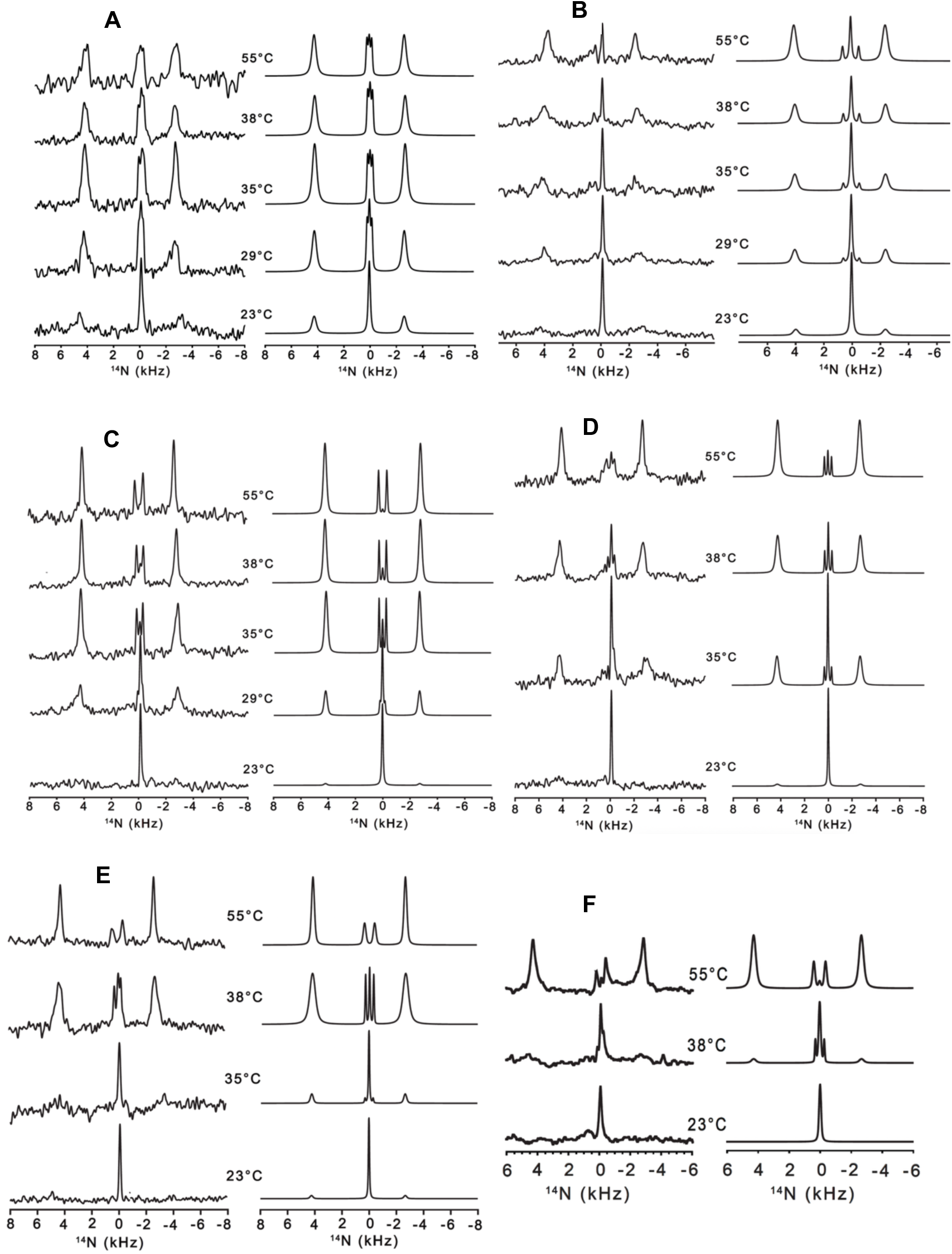
^14^N solid-state NMR spectra of DMPC (100 mg/mL) + 0.2 GA (20 mg/mL) bicelles recorded as a function of temperature (values indicated within each panel) in 10 mM Tris buffer containing 100 mM NaCl (pH 7.4) at varying cholesterol concentrations. Experimental spectra are shown on the left and the corresponding simulations on the right. The spectra were quantitatively fitted using a three-component bicelle model consisting of large aligned bicelles (B(L)), small bicelles (B(S)), and a residual isotropic/random component (B(R)). All components were modeled assuming an axially symmetric electric field gradient (η = 0). Cholesterol concentrations are (A) 0 mol%, (B) 5 mol%, (C) 10 mol%, (D) 15 mol%, (E) 20 mol%, and (F) 25 mol%. All additional simulation parameters for panels A–F are provided in Table TS2.

At 5 mol% cholesterol (Figure 3B & Table TS2), the increase in aligned bicelle population is small but steady. At 23 °C, isotropic or random assemblies dominate (B(R) = 74%), and only 26% are aligned bicelles. As temperature rises, B(L) gradually increases to 48% at 29 °C, 51% at 35 °C, and 61% at 38 °C, reaching 73% at 55 °C. The small bicelle population (B(S)) stays low (0–9%) at all temperatures. Compared to 0 mol%, the rise in B(L) occurs at higher temperatures, indicating that cholesterol stabilizes disordered or vesicular structures at low temperatures but still allows strong alignment at higher temperatures. This fits with cholesterol’s known role in making bilayers more ordered and rigid.^38–48^

At 10 mol% cholesterol (Figure 3C & Table TS2), the system behaves differently and more optimally. At 23 °C, it is almost entirely isotropic (B(R) = 92%), showing that aligned structures are suppressed at low temperature. But as temperature increases, there is a quick shift: B(L) jumps to 54% at 29 °C and reaches 72–79% between 35 and 55 °C, while B(R) drops sharply to 8% or less, eventually to about 1% at 55 °C. The small bicelle population (B(S)) increases modestly (7–20%), suggesting the presence of temporary intermediate structures during the transition. The aligned bicelle fraction stays high (about 79%) over a wide temperature range, showing stable and efficient alignment. This differs from both lower (0–5 mol%) and higher cholesterol systems and suggests that 10 mol% cholesterol provides the best balance between membrane fluidity and order. The quick drop in B(R) and steady B(L) support the idea that this composition is best for forming large, well-aligned bicelles.

At 15 mol% cholesterol (Figure 3D & Table TS2), the temperature-driven redistribution is broader and less cooperative. At 23 °C, isotropic assemblies dominate (B(R) = 87%), and there are few aligned bicelles (13%). As temperature rises, B(L) goes up to 62% at 35 °C and 77% at 38 °C, reaching 88% at 55 °C. However, compared to 10 mol%, the increase in B(L) happens over a wider temperature range, and significant isotropic and small bicelle populations remain at intermediate temperatures. This means that higher cholesterol stabilizes mixed phases and delays the formation of fully aligned bicelles.

At 20 mol% cholesterol (Figure 3E & Table TS2), this trend is even stronger. At 23 °C, the system is mostly isotropic (B(R) = 85%), with only 15% of the bicelles aligned. Even at 35 °C, isotropic components are still the majority (48%), and only at higher temperatures (38–55 °C) does B(L) rise to 79% and 77%. The continued presence of B(R) and the increase in B(S) (up to 23%) show more structural variety and less cooperative transition. These results suggest that higher cholesterol levels make the bilayer more rigid, thereby stabilizing disordered or partly ordered phases over a wider temperature range.

The 25 mol% cholesterol system (Figure 3F & Table TS2) shows the most extreme case. At 23 °C, the system is fully isotropic (B(R) = 100%), meaning bicelle formation is completely suppressed. At 38 °C, there is a significant range (B(L):B(S):B(R) = 18:22:60), indicating that all three components coexist. Only at 55 °C does the aligned bicelle population become dominant (B(L) = 78%), though a small amount of B(S) (20%) remains. This means that very high cholesterol leads to multiple bicelle environments and strong phase variety, which matches the need for multiple bicelle components seen in the ^31^P analysis. Across all cholesterol levels, the ^14^N results clearly show that cholesterol controls bicelle alignment by shifting phase populations rather than altering the basic headgroup structure. The steady values of δ_iso_ (about 27 ppm), C_q_ (about 8.5–9.2 kHz), and η = 0 confirm that the local electrostatic environment of the choline headgroup stays mostly the same. Instead, cholesterol changes the balance between isotropic assemblies, small bicelles, and large aligned bicelles. More cholesterol shifts the growth of B(L) to higher temperatures and widens the range of temperatures where B(S) and B(R) coexist, reflecting greater membrane rigidity and less cooperative alignment. Importantly, the B(L) populations measured by ^14^N NMR match well with the appearance and strength of the aligned bicelle peak in the ^31^P spectra, showing a direct link between local headgroup ordering and large-scale magnetic alignment. This connection confirms that the growth of large, well-ordered bicelles is the main factor for alignment efficiency.

Of all the compositions, 10 mol% cholesterol gives the best balance, with quick temperature-driven conversion to aligned bicelles, consistently high B(L) populations (about 70–80%), and very little isotropic content. This means that at this level, the membrane finds an ideal balance between fluidity and order, with sufficient flexibility for bicelle formation and reorganization but sufficient rigidity for stable alignment. As a result, large, well-ordered bicelles form efficiently over a wide temperature range, with little phase variation. These findings agree with earlier studies showing that moderate cholesterol levels improve membrane ordering and support stable bicelle alignment, while excessive cholesterol increases rigidity and disrupts alignment.^44–48,50,124–128^

### 3.4. Correlated ^31^P-^14^N NMR Reveals Cholesterol Dependent Bicelle Population Redistribution and Identifies an Optimal Alignment Regime

The combined ^31^P and ^14^N solid-state NMR data demonstrates the influence of cholesterol on bicelle formation and magnetic alignment for DMPC + 0.2 GA systems. The ^31^P spectra indicate global membrane alignment through a characteristic resonance near −10 ppm, while ^14^N NMR provides local structural information by quantifying lipid populations as large aligned bicelles (B(L)), small bicelles (B(S)), and isotropic or random assemblies (B(R)). These measurements establish a connection between overall alignment and molecular-level organization.^20,24,59,83,84,128^ Figure S9 presents heat maps of B(L) and B(R) that illustrate the effects of temperature and cholesterol on these populations. The B(L) heat map reveals that aligned bicelles increase with temperature across all samples, with the efficiency of this process dependent on cholesterol concentration. At 0 mol% cholesterol, B(L) increases gradually with temperature, whereas at 5 mol%, the transition is slower. At 10 mol% cholesterol, B(L) increases sharply and remains high (approximately 72–79%) over a broad temperature range (35–55 °C). In contrast, the B(R) heat map shows that isotropic populations decrease rapidly from about 92% at 23 °C to only 1–8% at elevated temperatures. At higher cholesterol concentrations (15–25 mol%), the increase in B(L) occurs at higher temperatures, and B(R) persists longer, indicating delayed bicelle formation and increased structural heterogeneity. These heat maps underscore the significant role of cholesterol in modulating the conversion of isotropic structures to aligned bicelles. Figure 4 further elucidates the transformation pathway by depicting the temperature-dependent changes in B(L), B(S), and B(R). In all samples, B(L) increases with temperature, but the transition dynamics differs. At low cholesterol (0–5 mol%), the transition is direct, with B(L) increasing smoothly and B(R) decreasing, while B(S) remains low. At higher cholesterol levels (15–25 mol%), a broader region emerges in which B(S) becomes more prominent, and B(R) decreases more gradually, indicating a less cooperative, more heterogeneous transition. At 10 mol% cholesterol, the system exhibits the highest efficiency: B(L) increases rapidly, B(R) decreases sharply, and B(S) remains moderate, reflecting a rapid, cooperative shift to aligned bicelles.

**Figure 4.**
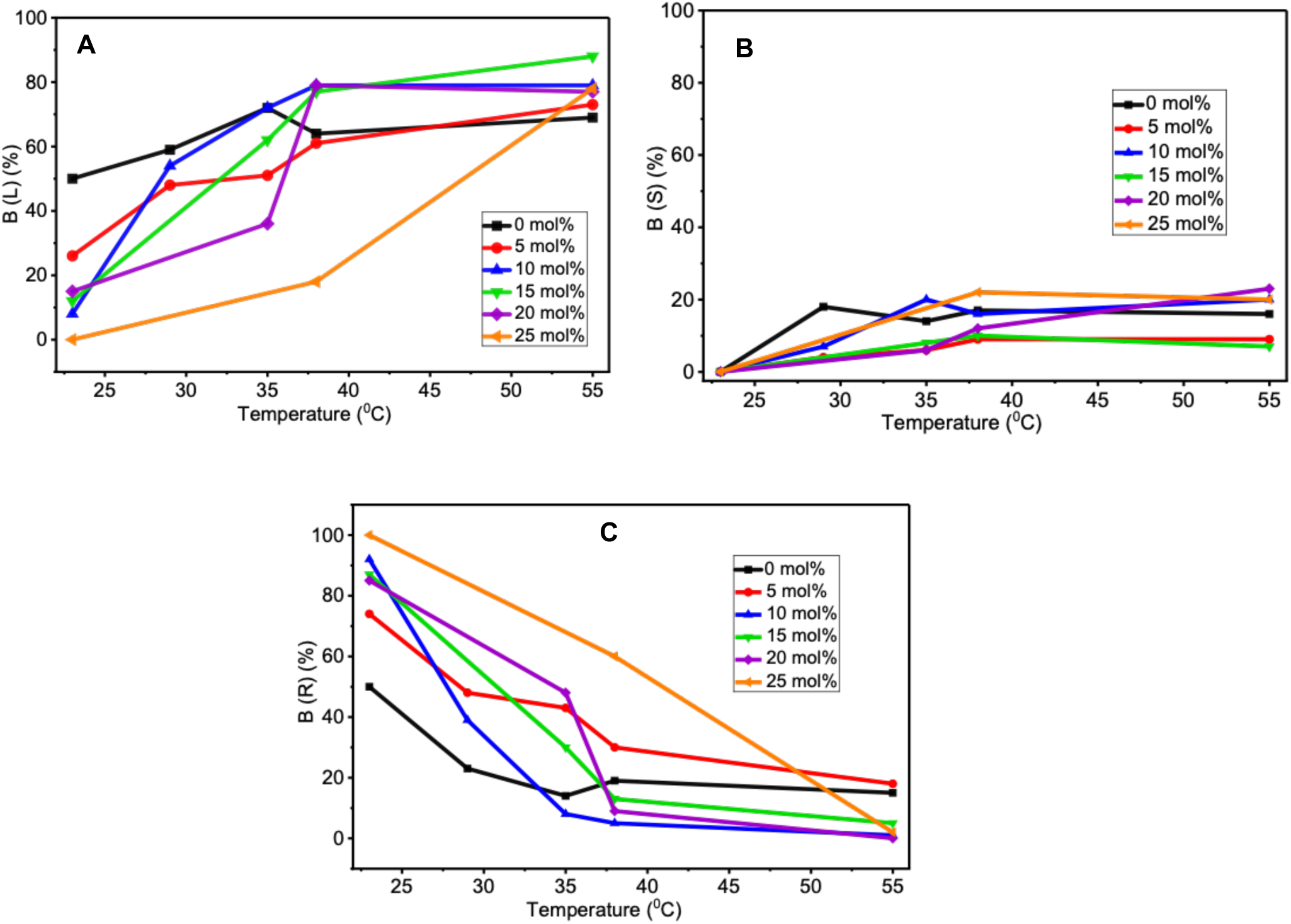
Cholesterol-dependent redistribution of bicelle populations showing (A) large aligned bicelles B(L), (B) small bicelles B(S), and (C) isotropic/random fraction B(R). The data illustrates a temperature-driven shift from isotropic assemblies to aligned bicelles, with B(R) dominating at low temperatures and progressively decreasing as B(L) increases upon heating. The transient presence of B(S) reflects intermediate bicelle states during structural reorganization. At ∼10 mol% cholesterol, the transition is most efficient, yielding a rapid increase in B(L) with minimal residual B(S) and B(R), whereas higher cholesterol concentrations (≥15 mol%) exhibit increased heterogeneity and delayed alignment, consistent with cholesterol-induced membrane ordering and domain formation.

## 4.0. Discussion

The present study provides a comprehensive mechanistic framework for understanding bicelle formation in DMPC + GA systems and the role of cholesterol in modulating membrane organization, as revealed by combined ^31^P and ^14^N solid-state NMR. Cholesterol’s effect on DMPC + GA bicelle formation observed here is consistent with its well-known condensing action on phosphatidylcholine bilayers, where it increases acyl-chain order and bilayer rigidity in a concentration-dependent manner.^124,125^ The progressive delay in the vesicle-to-bicelle transition with increasing cholesterol (0–25 mol%) mirrors reports that cholesterol increases bilayer bending rigidity, an effect confirmed by neutron spin-echo and ^2^H NMR studies showing up to a threefold rise in rigidity as cholesterol content increases.^125^

A key finding here is that this relationship is non-monotonic: 10 mol% cholesterol, not the cholesterol-free system or higher loadings, gives the fastest and most cooperative transition to a stable aligned bicelle phase. This agrees with the concept of a cholesterol "sweet spot," where moderate cholesterol optimizes membrane order without excessively restricting flexibility, while higher concentrations shift the system toward a more rigid, less dynamically responsive state.^124,126^ This pattern is also consistent with cholesterol-doped bicelle studies showing that moderate cholesterol improves magnetic alignability and bicelle disk size, whereas EPR studies of DHPC-containing bicelles show ordering rising steadily with cholesterol up to 20 mol%, without necessarily improving alignment efficiency at the highest levels tested.^38,117,127^

The complementary ^31^P and ^14^N data support this interpretation. ^31^P NMR is an established global probe of phospholipid alignment through its orientation-sensitive chemical shift anisotropy,^20^ while ^14^N NMR reports on the local electrostatic environment and motion of the choline headgroup, allowing detection of local ordering that precedes long-range alignment detectable by ^31^P NMR.^59^ The correspondence between the two datasets across all cholesterol levels indicates that local headgroup reorganization and global magnetic alignment are closely linked in this system.

Glycyrrhizic acid (GA) functioned here as an effective bicelle-forming amphiphile, consistent with recent reports that saponins, including glycyrrhizic acid, can solubilize DMPC into magnetically alignable bicelles.^30^ The temperature-driven vesicle-to-bicelle transition observed for the GA system is also consistent with related work on DMPC/glycyrrhizin mixtures showing a reversible, temperature-dependent conversion between vesicles and bicelle-like structures.^101^

Comparing this system to the well-studied DMPC/DHPC bicelle platform highlights both similarities and differences. In DHPC systems, alignment is governed by the long- to short-chain lipid ratio (q), with the short-chain lipid undergoing fast exchange between the planar bilayer and curved rim, a mechanism established through ^31^P NMR and the mixed bicelle model.^105^ Solution NMR studies of isotropic DMPC/DHPC bicelles have similarly shown that short-chain lipid dynamics and exchange govern lineshape averaging,^121^ and diffusion NMR studies of water and short-chain lipid mobility support this exchange-based model of bicelle organization.^123^ Cholesterol incorporation into DHPC-based bicelles has been shown to increase magnetic alignability and disk size,^38^ to broaden and thermally stabilize the alignment window when added as cholesterol sulfate,^52^ and to increase bicelle ordering progressively with concentration as measured by EPR.^117^ Our GA-based system reproduces this general pattern of cholesterol-modulated alignment, with the addition of a saponin, rather than a short-chain phospholipid, serving as the rim-stabilizing component.

Cholesterol is a near-universal component of eukaryotic membranes and plays a central role in organizing lipid rafts and liquid-ordered domains that influence membrane protein function.^2–5^ Consequently, bicelle systems used as membrane mimetics for structural biology benefit from incorporating physiologically relevant cholesterol while retaining the robust magnetic alignment required for oriented solid-state NMR.^9,20,24^ Bicelles enriched in cholesterol and sphingolipids have already been developed specifically to better approximate raft-like membrane environments for membrane protein structural studies,^43^ and cholesterol has more generally been used to modulate bicelle stability and composition for NMR-based structural applications.^50,53^ Solid-state NMR structure determination of membrane proteins in bicelles depends critically on achieving uniform, stable alignment over practical sample-handling temperatures,^58,75–79,129^ so the identification of an optimal cholesterol composition (10 mol%) that maximizes alignment efficiency in the DMPC + GA system provides a useful practical reference point for future bicelle designs intended to combine physiological cholesterol content with high-quality structural NMR data.

In summary, cholesterol modulates DMPC + GA bicelle formation and alignment in a concentration-dependent with 10 mol% cholesterol representing an optimal balance between membrane order and flexibility that supports rapid, stable bicelle alignment. These results extend established principles from cholesterol-membrane biophysics and the DMPC-DHPC bicelle studies to a saponin-stabilized bicelle system, offering practical guidance for future cholesterol-containing bicelle preparations for membrane protein structural studies.

## Acknowledgements

This work was funded in part by the NIH (R35GM13973 to AR). A portion of this work was performed at the National High Magnetic Field Laboratory, which is supported by National Science Foundation Cooperative Agreement No. DMR-2128556* and the State of Florida.

## Notes

### Competing Interest Statement

The authors have declared no competing interest.

