## Supporting Information for "Effects of Cholesterol on Nanodisc Formation and Magnetic Alignment in DMPC and Glycyrrhizic Acid Systems Probed by ^31^P and ^14^N Solid-State NMR"

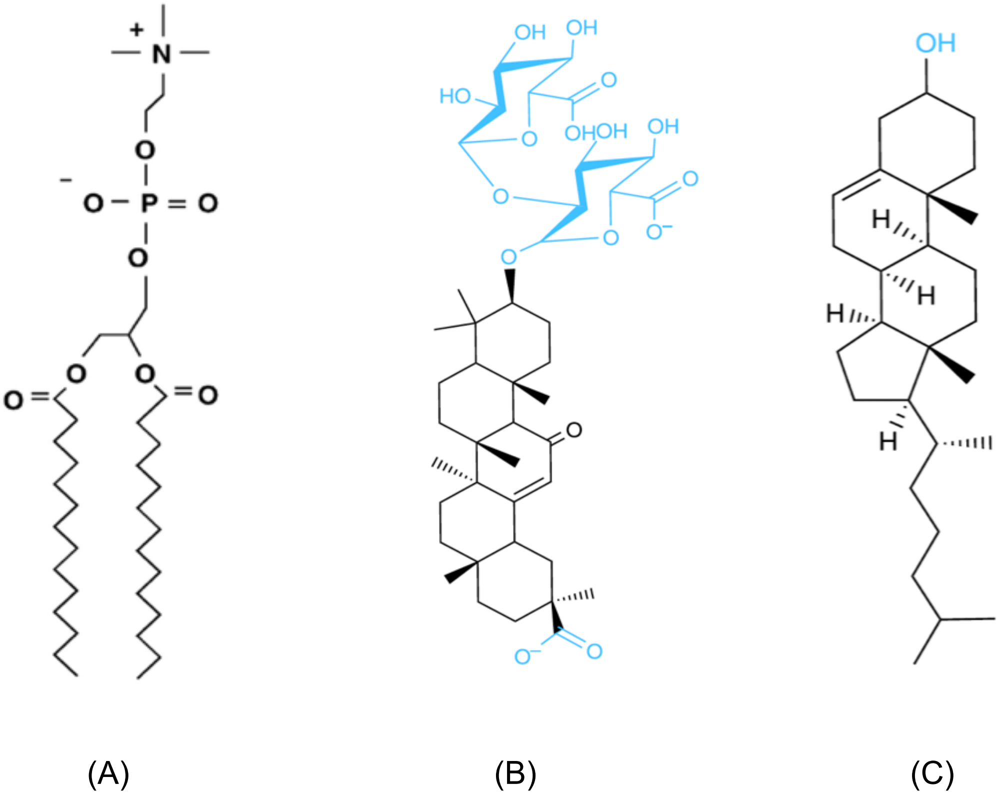


**Figure S1:** Molecular structures of (A) DMPC phospholipid, (B) glycyrrhizic acid (GA), and (C) cholesterol.

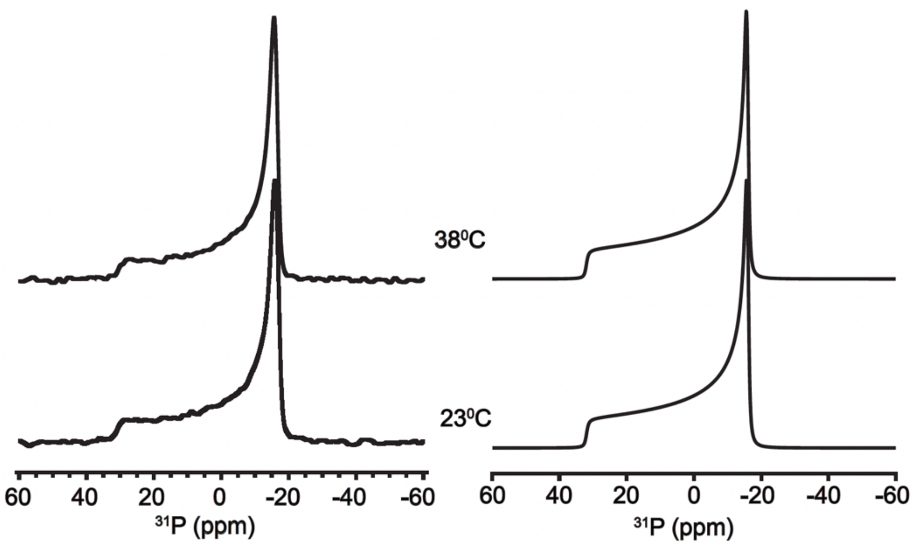


**Figure S2:** Experimental (left) and simulated (right) static $^{31}P$ solid-state NMR spectra of DMPC multilamellar vesicles (MLVs**)** acquired at 23 °C and 38 °C. The spectra exhibit the characteristic $^{31}P$ chemical shift anisotropy (CSA) powder pattern at both temperatures, consistent with phospholipid bilayers in the lamellar phase. The simulated spectra were generated using a CSA of 32 ppm and an axially symmetric CSA tensor ($\eta$ = 0), providing excellent agreement with the experimental lineshapes.

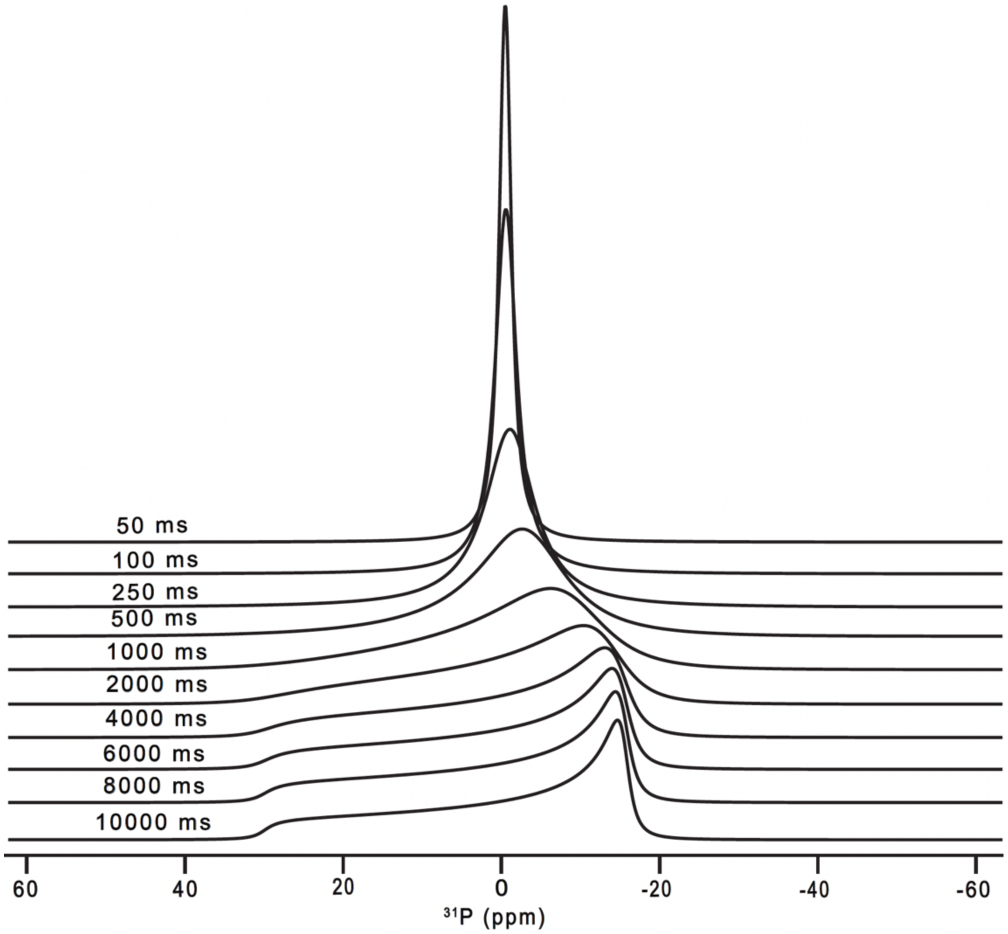


**Figure S3:** Simulated static $^{31}P$ solid-state NMR spectra showing the effect of molecular tumbling on the phosphorus chemical shift line shape. The spectra were simulated for correlation times ($\tau_c$) ranging from 10,000 ms (slow motion) to 50 ms (fast motion), corresponding to increasing molecular tumbling rates ($1/\tau_c$). Slow molecular motion preserves the full CSA powder pattern, whereas increasing tumbling progressively averages the anisotropic interaction, resulting in spectral narrowing and enhanced isotropic intensity. In the fast-motion limit ($\tau_C$ = 50 ms), complete motional averaging produces a sharp isotropic resonance at approximately 0 ppm.

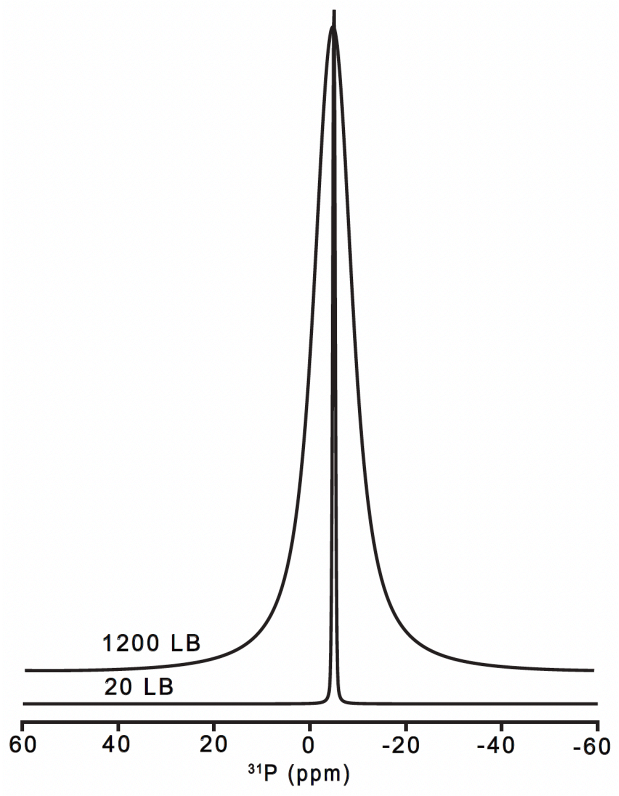


**Figure S4:** Simulated static $^{31}P$ solid-state NMR spectra illustrating the effect of Lorentzian line broadening (LB) on the spectral lineshape. The spectra were simulated using LB values of 20 and 1200 Hz, demonstrating that increasing line broadening significantly broadens the resonance and reduces spectral resolution while preserving the isotropic chemical shift near -5 ppm.

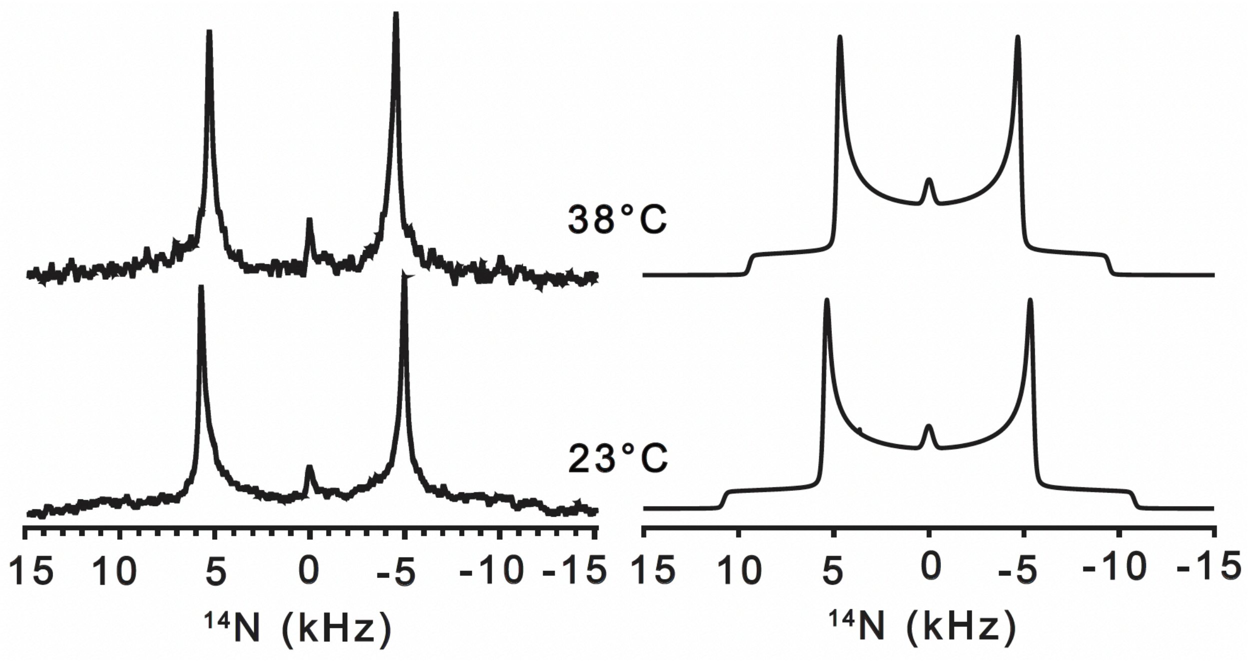


**Figure S5**: $^{14}$N solid-state NMR spectra of DMPC multilamellar vesicles (MLVs, 12% w/v) recorded in 10 mM Tris buffer containing 100 mM NaCl (pH 7.4). The left panels show the experimental spectra, while the right panel displays the corresponding simulated spectra.

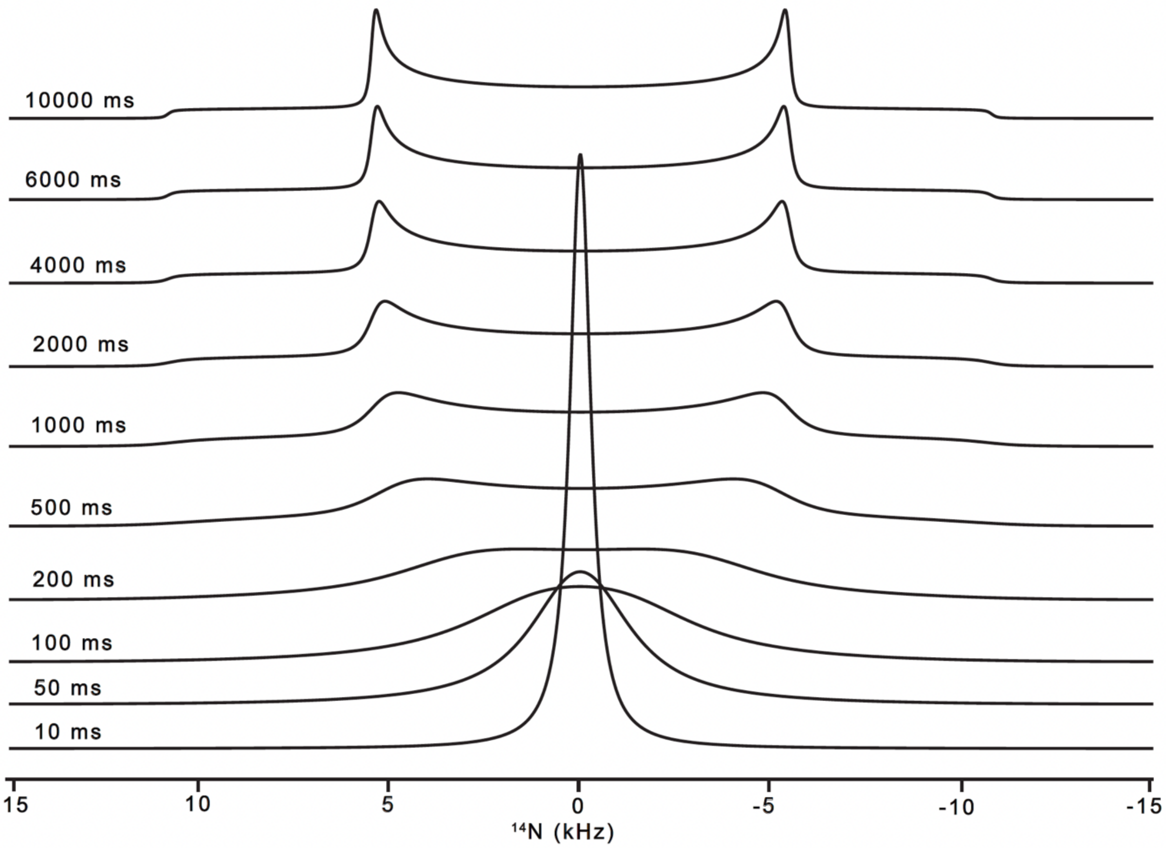


**Figure S6:** Simulated static $^{14}N$ solid-state NMR spectra showing the effect of molecular tumbling on the $^{14}N$ quadrupolar lineshape. The spectra were simulated for correlation times ($\tau_c$) from 10,000 ms (slow motion) to 10 ms (fast motion), corresponding to increasing tumbling rates ($1/\tau_c$). At 10,000–4000 ms, the spectra display the characteristic powder pattern of slowly tumbling phospholipid vesicles. As $\tau_c$ decreases (2000–200 ms), the powder pattern is progressively averaged, and at 100–10 ms, rapid tumbling produces a narrow isotropic resonance at 0 kHz, characteristic of fast-tumbling membrane assemblies such as small vesicles or micelles.

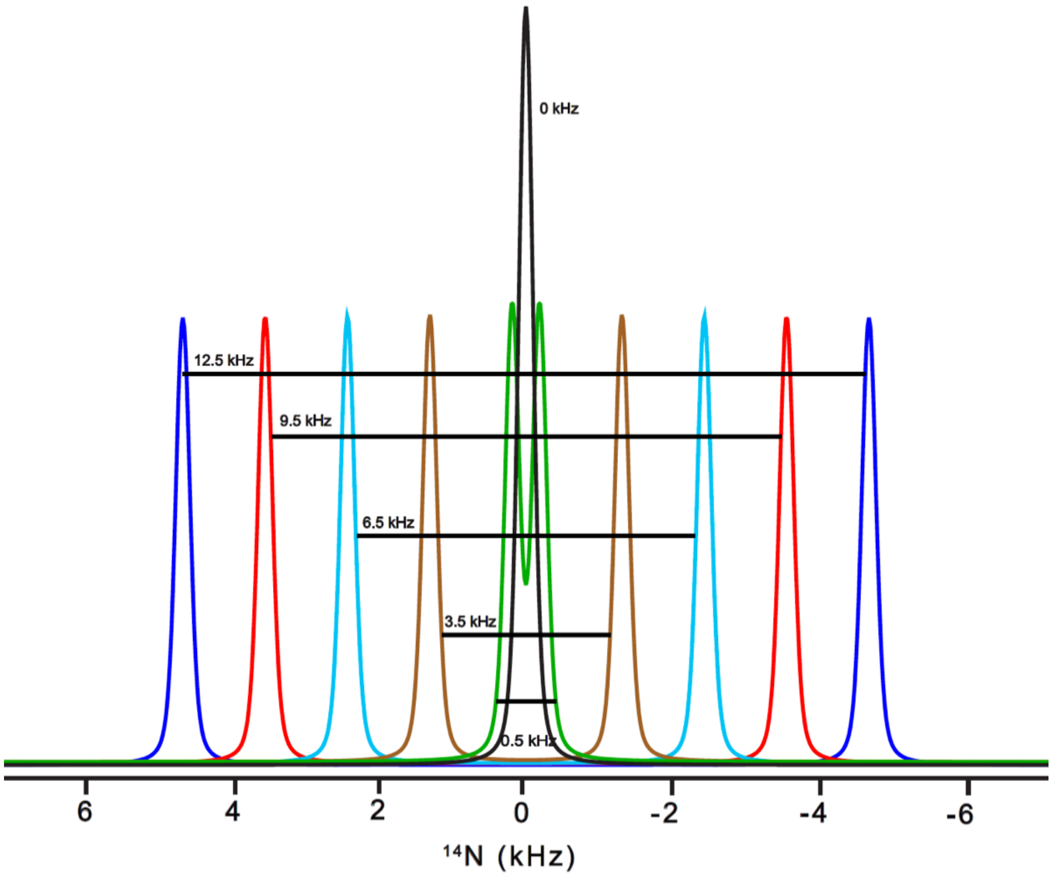


**Figure S7:** Simulated static $^{14}N$ solid-state NMR spectra showing the effect of increasing the quadrupolar coupling constant ($C_Q$) on the spectral lineshape. Simulations were performed for $C_Q$ values of 0.5, 3.5, 6.5, 9.5, and 12.5 kHz. Increasing $C_Q$ progressively enhances the quadrupolar splitting and broadens the spectral pattern, while decreasing $C_Q$ collapses the spectrum toward a single resonance at 0 kHz.

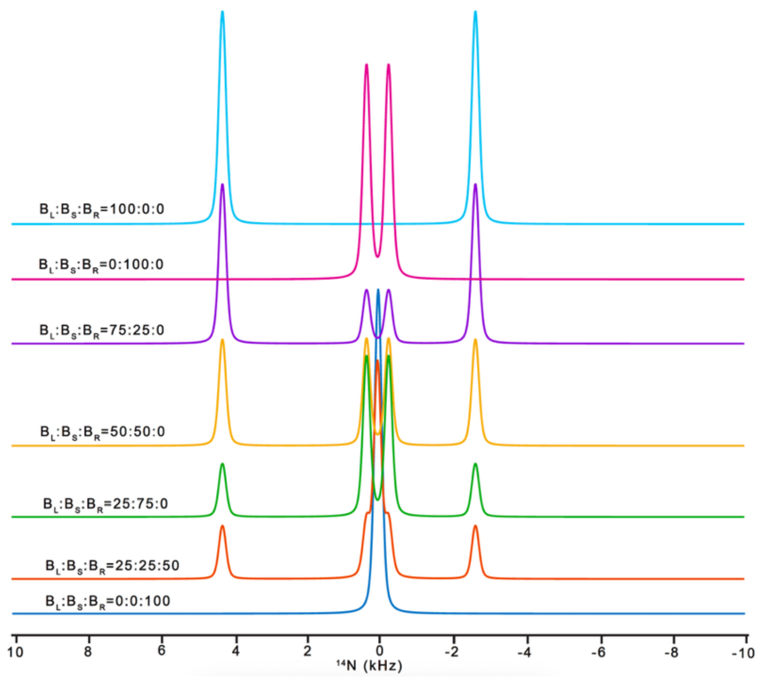


**(A)**

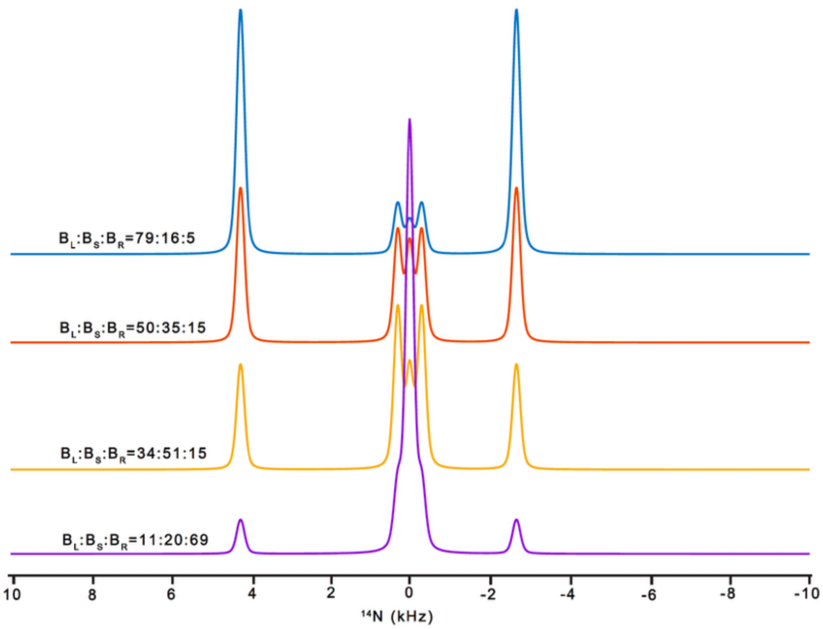


**Figure S8:** $^{14}$N NMR spectra of DMPC+0.2GA bicelle systems showing the evolution of lipid populations across different compositions. Spectra are plotted as a function of the relative populations of large bicelles ($B_L$), small bicelles ($B_S$), and random aggregated species including vesicles ($B_R$), with $B_L$:$B_S$:$B_R$ ratios indicated for each trace. The aligned Cont$^n$ figure S8: large bicelle component is centered near $\delta_{iso} \approx 27$ ppm with $C_q \approx 9.2$ kHz and axial symmetry ($\eta = 0$), while the small/isotropic bicelle component appears near $\delta_{iso}$ $\approx 0$ ppm with $C_Q \approx 0.5$ kHz. Top panel: progression from pure large bicelles (100:0:0, top (A)) to pure random species (0:0:100, bottom (B)). Bottom panel: selected intermediate compositions highlighting the coexistence of aligned and isotropic components.

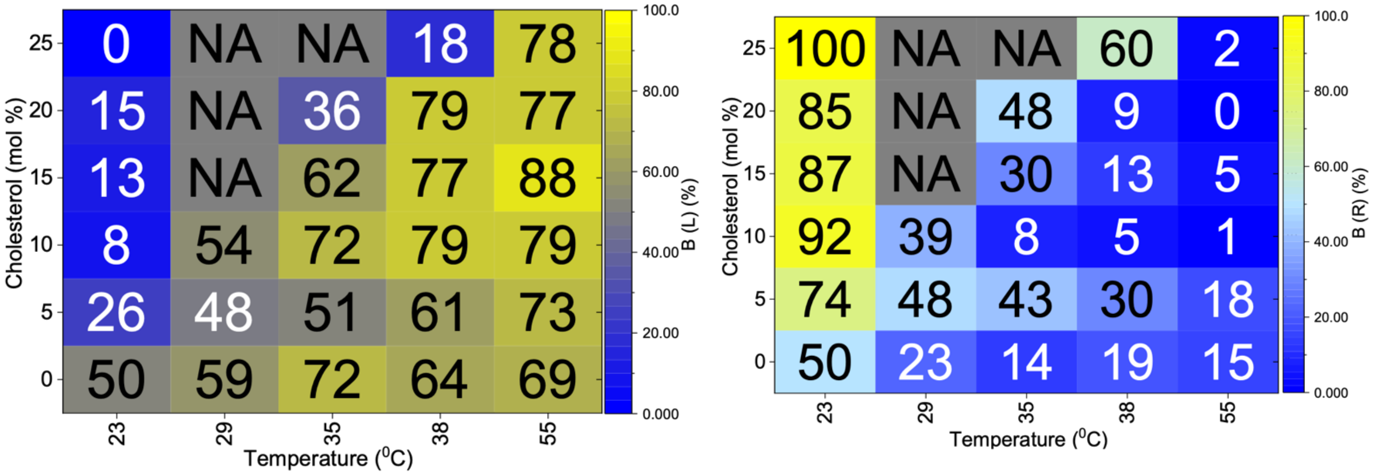


**(A)** **(B)**

**Figure S9:** Temperature-cholesterol heat map of (A) aligned bicelle fraction B(L) and (B) isotropic/random fraction B(R) obtained from $^{14}N$ solid-state NMR simulations.

**Table TS1:** Quantitative phase populations and simulation parameters extracted from static ³¹P solid-state NMR spectra of DMPC (100 mg/mL) + 0.2 GA (20 mg/mL) bicelles at varying cholesterol contents (0–25 mol%) and temperatures. Phase populations correspond to vesicles (V), aligned bicelles (B1), and, where required, a second bicelle population (B2). Simulations were performed assuming an axially symmetric CSA ($\eta = 0$) with a dominant value of ~20 ppm for all systems except 25 mol% cholesterol, where two bicelle environments (~20 and ~30 ppm) were required. Vesicle correlation times ($\tau_c$) reflect heterogeneous dynamics and decrease with increasing temperature, indicating enhanced lipid mobility. Line broadening (LB) captures increasing structural heterogeneity, particularly at higher cholesterol concentrations. The data demonstrate that increasing cholesterol delays bicelle formation, broadens phase coexistence, and introduces multiple bicelle environments at high sterol content.

| Chol (mol%) | Temp (°C) | Vesicles (V%) | | Bicelles (B%) | | τc (ms) | LB (Hz) |
|---|---|---|---|---|---|---|---|
| | | V1 | V2 | B1, CSA (20 ppm) | B2, CSA (30 ppm) | | |
| 0 | 23 | 100 | - | 0 | - | - | - |
| | 29 | 0 | - | 100 | - | - | - |
| | 32 | 0 | - | 100 | - | - | - |
| | 35 | 0 | - | 100 | - | - | - |
| | 38 | 0 | - | 100 | - | - | - |
| | 50 | 0 | - | 100 | - | - | 200 |
| | 55 | 0 | - | 100 | - | - | 800 |
| | 60 | 0 | - | 100 | - | - | 900 |
| 5 | 23 | 100 | - | 0 | - | - | - |
| | 29 | 85 | - | 15 | - | - | - |
| | 32 | 50 | - | 50 | - | - | - |
| | 35 | 22 | - | 78 | - | - | - |
| | 38 | 0 | - | 100 | - | - | 200 |
| | 55 | 0 | - | 100 | - | - | 400 |
| | 60 | 0 | - | 100 | - | - | 400 |
| 10 | 23 | 60 | 40 | 0 | - | 300/20000 | 200/500 |
| | 29 | 80 | 20 | 0 | - | 300/13000 | 200/500 |

| | | | | | | | |
|---|---|---|---|---|---|---|---|
| | 32 | 50 | - | 50 | - | 8000 | 600/500 |
| | 35 | 40 | - | 60 | - | 7000 | 600/700 |
| | 38 | 40 | - | 60 | - | 2000 | 600/900 |
| | 55 | 40 | - | 60 | - | 1300 | 600/900 |
| | 60 | 40 | - | 60 | - | 1300 | 600/900 |
| 15 | 23 | 80 | 20 | 0 | - | 200/4000 | 200/500 |
| | 32 | 35 | 65 | 0 | | 200/4000 | 200/500 |
| | 35 | 10 | 90 | 0 | - | 200/4000 | 400/500 |
| | 38 | 5 | 95 | 0 | - | 200/4000 | 400/500 |
| | 46 | - | 95 | 5 | - | 4000 | 200/300 |
| | 55 | 10 | - | 90 | - | 4000 | 200/600 |
| | 60 | 5 | - | 95 | - | 4000 | 200/900 |
| 20 | 23 | 40 | 60 | 0 | - | 200/4000 | 200/500 |
| | 35 | 35 | 65 | 0 | - | 200/4000 | 200/500 |
| | 38 | 75 | - | 25 | - | 4000 | 500/800 |
| | 46 | 60 | - | 40 | - | 4000 | 500/800 |
| | 55 | 20 | - | 80 | - | 4000 | 500/800 |
| | 60 | 10 | - | 90 | - | 4000 | 500/800 |
| 25 | 23 | 90 | - | - | 10 | 50 | 70/100 |
| | 38 | 79 | - | 5 | 16 | 140 | 300/400/100 |
| | 55 | 50 | - | 40 | 10 | 140 | 300/400/100 |
| | 60 | 16 | - | 82 | 2 | 140 | 300/400/100 |

**Table TS2.** Quantitative phase populations and simulation parameters extracted from $^{14}N$ solid-state NMR spectra of DMPC (100 mg/mL) + 0.2 GA (20 mg/mL) bicelles at varying cholesterol contents and temperatures. Spectra were fitted using a three-component model consisting of large aligned bicelles B(L), small bicelles B(S), and isotropic/random assemblies B(R), assuming an axially symmetric electric field gradient ($\eta$ = 0) for all components. Across all compositions, the aligned bicelle component is centered near $\delta_{iso}$ ≈ 27 ppm with $C_q$ ≈ 8.5–9.2 kHz, whereas the small bicelle component appears near $\delta_{iso}$ ≈ 0 ppm with $C_q$ ≈ 0.5–1.5 kHz, reflecting stronger motional averaging. Increasing cholesterol shifts the increase in B(L) to higher temperature and broadens the coexistence of B(S) and B(R), indicating delayed alignment and enhanced structural heterogeneity. Values are compiled from the fitted population trends described in the 3.3 results text.

| Chol (mol%) | Temp (°C) | B(L) (%) | B(S) (%) | B(R) (%) | B(L): $\delta_{iso}/C_q$ (ppm/kHz) | B(S): $\delta_{iso}/C_q$ (ppm/kHz) |
|---|---|---|---|---|---|---|
| 0 | 23 | 50 | 0 | 50 | 27 / 9.2 | 0 / 0.5 |
| | 29 | 59 | 18 | 23 | 27 / 9.2 | 0 / 0.5 |
| | 35 | 72 | 14 | 14 | 27 / 9.2 | 0 / 0.5 |
| | 38 | 64 | 17 | 19 | 27 / 9.2 | 0 / 0.5 |
| | 55 | 69 | 16 | 15 | 27 / 9.2 | 0 / 0.5 |
| 5 | 23 | 26 | 0 | 74 | 27 / 8.5 | 0 / 1.5 |
| | 29 | 48 | 4 | 48 | 27 / 8.5 | 0 / 1.5 |
| | 35 | 51 | 6 | 43 | 27 / 8.5 | 0 / 1.5 |
| | 38 | 61 | 9 | 30 | 27 / 8.5 | 0 / 1.5 |
| | 55 | 73 | 9 | 18 | 27 / 8.5 | 0 / 1.5 |
| 10 | 23 | 8 | 0 | 92 | 27 / 9.2 | 0 / 0.8 |
| | 29 | 54 | 7 | 39 | 27 / 9.2 | 0 / 0.8 |
| | 35 | 72 | 20 | 8 | 27 / 9.2 | 0 / 0.8 |
| | 38 | 79 | 16 | 5 | 27 / 9.2 | 0 / 0.8 |
| | 55 | 79 | 20 | 1 | 27 / 9.2 | 0 / 0.8 |
| 15 | 23 | 13 | 0 | 87 | 27 / 9.2 | 0 / 0.8 |
| | 35 | 62 | 8 | 30 | 27 / 9.2 | 0 / 0.8 |
| | 38 | 77 | 10 | 13 | 27 / 9.2 | 0 / 0.8 |
| | 55 | 88 | 7 | 5 | 27 / 9.2 | 0 / 0.8 |
| 20 | 23 | 15 | 0 | 85 | 27 / 9.2 | 0 / 0.8 |

| | | | | | | |
|---|---|---|---|---|---|---|
| | 35 | 36 | 6 | 48 | 27 / 9.2 | 0 / 0.8 |
| | 38 | 79 | 12 | 9 | 27 / 9.2 | 0 / 0.8 |
| | 55 | 77 | 23 | 0 | 27 / 9.2 | 0 / 1.0 |
| 25 | 23 | 0 | 0 | 100 | 27 / 9.2 | 0 / 1.0 |
| | 38 | 18 | 22 | 60 | 27 / 9.2 | 0 / 1.0 |
| | 55 | 78 | 20 | 2 | 27 / 9.2 | 0 / 1.0 |
